# Green-Synthesized Silver Nanoparticles from Coconut: A Novel Antimicrobial Strategy Against Acne Pathogens

**DOI:** 10.1101/2025.09.02.673756

**Authors:** Risav Banerjee, Kumkum Nag, Sneha Deb, V Suneetha

## Abstract

Acne vulgaris is a widespread dermatological disorder, often complicated by multidrug-resistant (MDR) bacteria such as *Staphylococcus aureus*. Rising resistance to conventional antibiotics has created an urgent need for alternative antimicrobial strategies. Nanoparticles, due to their nanoscale dimensions and unique surface properties, have shown considerable promise in overcoming bacterial resistance mechanisms.

In this study, acne-associated bacteria were isolated and identified through biochemical and microscopic methods, followed by antibiotic susceptibility testing. The isolates exhibited resistance to commonly prescribed antibiotics including vancomycin, erythromycin, ampicillin, and amoxicillin. To address this, silver nanoparticles (AgNPs) were synthesized using coconut shell extract (CSE), a waste-derived, phenolic-rich material that acted both as a reducing and stabilizing agent in the green synthesis process.

CSE-AgNPs demonstrated strong antibacterial activity against MDR *S. aureus*, producing clear zones of inhibition, reduced MIC values, and structural damage to bacterial cell walls. These findings highlight CSE-AgNPs as an eco-friendly, cost-effective, and potent antimicrobial alternative, integrating waste valorization with nanotechnology to combat antibiotic resistance in acne-associated pathogens.

## Introduction

Acne vulgaris is a chronic inflammatory disorder of the pilosebaceous unit, encompassing the hair follicle, sebaceous gland, and associated structures^1^. It represents one of the most common dermatological conditions globally, affecting nearly 9–10% of the population and ranking among the leading causes of skin-related disability^2^. A meta-analysis of 42 studies confirmed a robust association between acne and elevated levels of depression (r = 0.22) and anxiety (r = 0.25)^3^. While often regarded as a benign condition of adolescence, severe or untreated acne can result in permanent scarring, secondary infections, and considerable psychological morbidity, including anxiety and depression^4^. The onset typically occurs between 12 and 14 years of age, with earlier manifestation in females, although persistence into adulthood is increasingly recognized^5^.

The clinical spectrum of acne is diverse and broadly categorized into three types:

- Mild acne is characterized by the presence of open and closed comedones (blackheads and whiteheads) with occasional papules. These lesions are often localized, most prominently on the nose, forehead, and chin^6^.
- Moderate acne is characterized by approximately 6 to 20 inflammatory lesions— namely papules and pustules—on half of the face, reflecting a marked escalation in lesion count and inflammation relative to mild forms^7^.
- Severe acne presents with 21 to 50 inflammatory eruptions (papules, pustules, and sometimes nodules) on half of the face, indicating extensive inflammation and higher risk of scarring and treatment resistance^8^.

The etiology of acne involves a multifactorial process, including increased sebum production, follicular hyperkeratinization, bacterial colonization, and heightened inflammatory responses^9^. The skin microbiota, particularly Gram-positive organisms such as *Staphylococcus* spp., *Micrococcus* spp., *Corynebacterium* spp., and *Cutibacterium acnes* (formerly *Propionibacterium acnes*), play a central role in pathogenesis^10^. Among these, *C. acnes* has long been recognized as a primary etiological agent, but recent studies have also implicated *Staphylococcus aureus* as a major contributor, particularly in moderate to severe and treatment-refractory cases^11^.

Of concern, acne-associated *S. aureus* isolates frequently display multidrug resistance (MDR), rendering them unresponsive to first-line antibiotics such as tetracyclines, macrolides (erythromycin, clindamycin), and even glycopeptides like vancomycin^12^. The rise of methicillin-resistant *S. aureus* (MRSA) and vancomycin-resistant *S. aureus* (VRSA) has further exacerbated therapeutic challenges, raising alarm among clinicians and researchers alike^13^.

To address these limitations, nanotechnology has emerged as a promising frontier in antimicrobial therapy. Metallic nanoparticles, particularly silver nanoparticles (AgNPs), are widely studied for their broad-spectrum antibacterial activity, which stems from multiple mechanisms including disruption of cell membranes, induction of oxidative stress^15^, and interference with DNA replication^16^. Unlike conventional antibiotics, these multitargeted effects reduce the likelihood of resistance development. Importantly, “green synthesis” of AgNPs using plant extracts offers a sustainable, low-cost, and environmentally benign method^17^, as phytochemicals such as phenols and flavonoids act as natural reducing and stabilizing agents^18^.

Coconut (*Cocos nucifera*) shells^19^, an abundant agricultural by-product, are rich in bioactive compounds including polyphenols, flavonoids (e.g., luteolin), and tannins with well-documented antioxidant and antimicrobial activities^18^. These phytochemicals facilitate the bioreduction of silver ions into stable AgNPs and enhance their antimicrobial potential through synergistic bioactivity^20^. Valorization of coconut shell waste^16^ for nanoparticle synthesis thus integrates waste recycling with biomedical innovation, offering both environmental and therapeutic benefits.

In this context, the present study focuses on the green synthesis of AgNPs using coconut shell extract (CSE) and their evaluation against MDR *S. aureus* isolated from acne patients. By characterizing the growth patterns, resistance profiles, and antimicrobial susceptibility of these isolates, we demonstrate the potential of CSE-AgNPs as an effective, eco-friendly alternative to conventional antibiotics for acne management.

Although numerous studies have investigated the antimicrobial potential of silver nanoparticles, the majority rely on chemical synthesis methods that require hazardous reducing agents, involve high costs, and raise environmental safety concerns^21^. While plant-mediated “green” synthesis approaches have been reported^20^, only a limited number of studies have directly explored the use of agro-waste materials, such as coconut shells, as both reducing and stabilizing agents^22^. Furthermore, many earlier investigations have tested AgNPs against common pathogens in general, without specifically addressing acne-associated multidrug-resistant *S. aureus*, which remains a growing clinical challenge^23^.

Another critical gap lies in the lack of comparative analyses between nanoparticle efficacy and conventional antibiotics in acne-related contexts. Previous reports have often demonstrated antibacterial activity qualitatively, but few have rigorously compared the zones of inhibition, MIC values, or morphological changes against MDR acne pathogens alongside frontline antibiotics^24^. Moreover, despite coconut-derived products being widely recognized for their bioactivity, the mechanistic contributions of coconut shell phytochemicals— particularly phenolics and flavonoids—in nanoparticle synthesis and antimicrobial activity have not been systematically explored.

The present study addresses these gaps by:

1. Utilizing coconut shell extract, a readily available agricultural waste, for the eco-friendly synthesis of AgNPs.
2. Testing the antibacterial efficacy of CSE-AgNPs specifically against MDR *S. aureus* isolated from acne patients.
3. Conducting comparative evaluations with commonly used antibiotics to highlight their therapeutic potential.

By bridging these gaps, our work not only provides a novel antimicrobial strategy but also underscores the potential of waste valorization in developing sustainable, cost-effective nanomedicines.

## Methods and Materials

All clinical material was processed in a Class II cabinet using BSL-2 practices. Slides and disposable materials were decontaminated by autoclaving or according to institutional biohazard waste protocols.

### Isolation of Bacterial Strains from Acne Lesions

Clinical samples were obtained from a patient presenting with mild adult acne within the age group of 20–25 years. The subject had no prior history of acne treatment, including topical or systemic retinoids (e.g., vitamin A derivatives), or antibiotic therapy such as lincosamides (clindamycin), tetracyclines (minocycline, doxycycline), or macrolides (erythromycin, azithromycin), ensuring the absence of antimicrobial interference. Following ethical approval and informed consent, acne lesions were carefully sampled under aseptic conditions. Blood and purulent material were aspirated from the inflamed lesion using a sterile syringe. The collected specimens were immediately transferred into sterile containers and processed under a laminar airflow cabinet to minimize contamination.

For bacterial isolation, aliquots of the samples were inoculated onto selective and non-selective culture media, including nutrient agar, blood agar, and MacConkey agar plates, and incubated at 37 °C for 24–48 hours under both aerobic and anaerobic conditions to allow the growth of potential acne-associated microorganisms. Emerging colonies were sub-cultured to obtain pure isolates, which were subsequently subjected to standard microbiological characterization.

### Media Preparation and Culture Conditions

Nutrient agar medium was prepared by dissolving 1.3 g of nutrient agar powder in 100 mL of distilled water. The suspension was brought to boiling to ensure complete dissolution and subsequently sterilized in an autoclave at 121 °C for 15 minutes at 15 lbs pressure. Sterile Petri plates were poured inside a laminar airflow chamber and allowed to solidify under aseptic conditions. Plates were incubated at 37 °C for 24 hours to confirm sterility.

Clinical samples, consisting of pus admixed with traces of blood, were obtained from acne lesions using sterile swabs. These were immediately inoculated onto freshly prepared nutrient agar plates. The four-quadrant streaking method was applied to dilute the inoculum progressively, enabling separation of bacterial colonies from mixed populations. The plates were incubated at 37 °C for 24 hours, after which discrete colonies appeared. Well-isolated colonies were sub-cultured repeatedly to obtain pure cultures for downstream analyses.

### Morphological and Microscopic Characterization

#### Gram Staining

Gram staining was performed to classify clinical and culture isolates by cell wall architecture and to document cellular morphology and arrangement. The procedure followed a standard crystal-violet/iodine–ethanol–safranin method with defined timings and in-run quality controls.

From a fresh (18–24 h) colony grown on nutrient agar, a small amount was emulsified in 10– 20 µL sterile saline on a clean slide to form a thin, even smear (∼1 cm diameter). After that, it was made air-dry completely (≥5 min). For heat fixation, the fully dried smear was made pass through a low flame 2–3 times, smear side up (1–2 s per pass).

#### Staining procedure

Primary stain (crystal violet): The smear was flooded for 60 s. It was rinsed gently with distilled water while holding the slide at ∼45° to prevent reagent pooling.

Mordant (Gram’s iodine): Again, it was flooded for 60 s to form the crystal-violet–iodine complex, then it was rinsed with water.

#### Decolorization

With the slide tilted at 45°, 95% ethanol was poured dropwise over the smear for approximately 10–20 s or until the runoff is almost colourless, moving the stream across the smear. Then again it was rinsed with water.

#### Counterstain (safranin)

Lastly, it was flooded with Safranin for 30–60 s and again rinsed with water and gently blotted dry with lint-free tissue. Also, it was air-dried briefly before microscopy.

#### Microscopy and image capture

It was examined with a brightfield microscope using 100× oil-immersion objective (total magnification 1000×). For each isolate, ≥30 non-overlapping fields were evaluated.

Gram reaction, cell shape (cocci/rods/pleomorphic), size (in µm) arrangement (clusters, chains, pairs, palisades), and background matrix (e.g., host cells, debris) was documented. Representative fields were imaged with a digital camera; scale bars were generated using a stage micrometer calibration. As an experimental control, smears of *S. aureus* ATCC 25923 and *E. coli* ATCC 25922 were used alongside test slides. *S. aureus* = uniformly purple cocci in clusters; *E. coli* = uniformly pink rods.

#### Capsule staining

Capsule staining was performed to detect extracellular polysaccharide or polypeptide capsules associated with virulence and biofilm formation.

#### Negative stain (India ink / nigrosin)

A single small drop (∼5–10 µL) of India ink was put near one end of a clean slide. Using a sterile loop, a small amount of a well-isolated, fresh colony was transferred into the drop and mixed gently to form a thin suspension. With a second slide holding at a low angle (≈30°), the drop was touched and the suspension was pulled along the slide to produce a thin smear approximately 1 cm long. The smear was allowed to air dry completely at room temperature. Lastly it was examined under brightfield microscopy using the 100× oil-immersion objective. Multiple non-overlapping fields (≥30) was observed.

#### Motility test — Hanging drop method

The hanging drop preparation was used to assess true bacterial motility (flagella- or gliding-mediated) and to distinguish it from Brownian motion or convective currents.

Using a sterile loop, a small amount of growth from a well-isolated, fresh colony (18–24 h) was picked and suspended in 50–100 µL sterile saline to produce a lightly turbid suspension. Then it was mixed gently to disperse clumps. The drop was directly placed in the cavity of the depression slide. The coverslip was inverted over the cavity slide so the drop hangs in the depression.

The slide was within 10–30 minutes under phase-contrast microscope. Multiple non-overlapping fields (at least 5–10 fields or ≥100 cells in total) were analysed to obtain a representative assessment.

### Biochemical Characterization of Isolates

A series of classical biochemical tests were performed to characterize the metabolic and enzymatic properties of the bacterial isolates. Standard protocols were followed with appropriate positive and negative controls to ensure accuracy and reproducibility.

#### Catalase Test

The catalase test was performed to detect the presence of the catalase enzyme, which decomposes hydrogen peroxide (H2O2) into water and oxygen, thereby protecting cells from oxidative damage.

Freshly grown colonies (18–24 h old) were transferred onto a clean glass slide using a sterile inoculating loop. A drop of 3% H2O2 was added directly to the colony. The immediate formation of visible effervescence (oxygen bubbles) was interpreted as a positive reaction, whereas the absence of bubbling indicated catalase negativity. *Staphylococcus aureus* served as a positive control and *Enterococcus faecalis* as a negative control.

#### Oxidase Test

The oxidase test was performed to assess the presence of cytochrome c oxidase, an enzyme involved in the electron transport chain.

Colonies (18–24 h old) were smeared onto oxidase reagent strips impregnated with *p*-phenylenediamine dihydrochloride. Development of a deep purple coloration within 10–30 seconds indicated a positive oxidase reaction, while no colour change (or change beyond 60 s) was considered negative. *Pseudomonas aeruginosa* was used as a positive control and *Escherichia coli* as a negative control. IMViC Test

The IMViC test (Indole, Methyl Red, Voges–Proskauer, Citrate) was used to differentiate members of the Enterobacterales and to provide further taxonomic resolution of the isolates.

1. Indole Test: Cultures were inoculated into tryptone broth and incubated at 37 °C for 24–48 h. Following incubation, 0.5 mL of Kovac’s reagent (isoamyl alcohol, *p*-dimethylaminobenzaldehyde, HCl) was added to each tube. Development of a distinct red ring at the liquid–air interface indicated indole production from tryptophan metabolism, confirming a positive reaction. Yellow or no color change was considered negative.
2. Methyl Red (MR) Test: Isolates were grown in MR-VP broth at 37 °C for 48 h. After incubation, five drops of methyl red indicator were added. A stable red coloration signified mixed acid fermentation and was recorded as a positive result. A yellow/orange color indicated a negative result.
3. Voges–Proskauer (VP) Test: Cultures grown in MR-VP broth were treated with 0.6 mL of α-naphthol followed by 0.2 mL of 40% KOH. Tubes were shaken gently and observed for 15–30 min. The appearance of a red coloration confirmed acetoin (neutral end-product) production and was scored as positive, whereas a yellow/brown reaction was considered negative.
4. Citrate Utilization Test: Isolates were streak-inoculated onto Simmons citrate agar slants and incubated at 37 °C for up to 48 h. Growth on the medium accompanied by a color change from green to blue indicated utilization of citrate as the sole carbon source, confirming a positive result. No growth or retention of the green color was interpreted as negative.

#### Triple Sugar Iron (TSI) Agar Test

To evaluate carbohydrate fermentation and hydrogen sulfide production, isolates were inoculated by stabbing the butt and streaking the slant of TSI agar. Following 24 h incubation at 37 °C, reactions were interpreted as follows:

- Acid production from glucose, lactose, and/or sucrose fermentation was indicated by a yellow color in the slant and/or butt.
- Gas production was identified by cracks, bubbles, or displacement of the agar.
- Hydrogen sulfide (H2S) production was evidenced by blackening of the butt due to ferrous sulfide precipitation.
- A red slant/yellow butt pattern suggested glucose fermentation only, while a completely red slant/butt indicated no carbohydrate utilization (alkaline reaction).

#### Mannitol Salt Agar (MSA) Test

Mannitol salt agar was employed as a selective and differential medium for the detection of *Staphylococcus aureus*. Isolates were streaked onto MSA plates containing 7.5% NaCl, which selectively supports staphylococcal growth due to their halotolerance. Fermentation of mannitol by *S. aureus* was indicated by a color change of the medium from red (alkaline) to bright yellow (acidic), as a result of phenol red indicator response to acid end-products. Non-mannitol fermenting staphylococci (e.g., *S. epidermidis*) grew on the medium but did not alter the color, thus remaining red. Absence of growth indicated salt sensitivity and exclusion from the genus *Staphylococcus*.

Collectively, these biochemical assays provided an integrated phenotypic profile of the clinical isolates, enabling presumptive identification of *Staphylococcus aureus* and other bacteria associated with acne lesions. The combination of enzymatic, metabolic, and selective/differential media-based assays increased the reliability of bacterial characterization in the absence of molecular confirmation such as 16S rRNA sequencing.

### Preparation of coconut-shell extract for AgNP synthesis

Collection and pre-processing: Mature coconut shells were obtained from a local market (Katpadi,Vellore, Tamil Nadu) cleaned thoroughly to remove adhering dust and foreign matter, and air-dried at room temperature. The dried shells were further oven-dried at 40–50 °C to constant weight, pulverized using a sterile mortar and pestle and passed through a 60-mesh sieve to obtain a uniform fine powder. The powdered material was stored in a desiccator in amber vials until extraction.

#### Weighing and solvent selection

A total of 2.685 g of the powdered coconut shell was weighed and divided into three equal portions (≈0.895 g each) for extraction with solvents of increasing polarity: diethyl ether (low polarity), *n*-butanol (intermediate polarity) and isopropanol (isopropyl alcohol; moderately polar). The solvent series was chosen to fractionate phytochemicals across a polarity range — nonpolar lipids/terpenoids in ether, medium-polarity phenolics and some flavonoids in butanol, and more polar polyphenols/reducing sugars in isopropanol — thereby maximizing recovery of potential reducing and capping agents for AgNP formation.

#### Extraction (maceration with agitation)

Each powdered aliquot (∼0.895 g) was extracted in 10 mL of the respective solvent (solid:solvent ratio ≈1:11 w/v). The mixtures were placed in sterile, amber screw-cap tubes and incubated on an orbital shaker at 150 rpm in a temperature-controlled incubator set to 35–37 °C for 24 h to facilitate solute diffusion. Gentle agitation and mild elevated temperature increase extraction efficiency while minimizing thermal degradation of labile phytochemicals.

#### Clarification and concentration

After incubation, the solvent extracts were clarified by gravity filtration through Whatman No. 1 filter paper followed by centrifugation at 5,000 × g for 10 min to remove fine particulates. Filtrates were combined where appropriate and concentrated to dryness using a rotary evaporator under reduced pressure (bath temperature 35–40 °C) to remove organic solvents. Residual solvent traces were removed by drying under a gentle stream of nitrogen or in a vacuum desiccator. All solvent handling and evaporation steps were performed in a fume hood and with appropriate safety precautions because diethyl ether and isopropanol are highly flammable.

#### Reconstitution

The dried extracts were reconstituted in deionized distilled water to prepare aqueous stock solutions. Reconstitution was performed with sonication (10 min, room temperature) and brief warming (≤40 °C) if necessary to fully dissolve water-soluble fractions. Because some nonpolar constituents may not be water-soluble, extracts were allowed to form colloidal suspensions when appropriate; these suspensions were homogenized by vortexing immediately prior to use. Typical working stock concentrations were prepared at 50–100 mg/mL (w/v) based on dry extract weight; the exact concentration used for nanoparticle synthesis and for antimicrobial assays was recorded for each batch.

#### Sterilization and storage

Aqueous extracts intended for microbiological testing were sterile-filtered through 0.22 µm syringe filters into sterile amber vials under aseptic conditions.

Filtered extracts were aliquoted to avoid freeze–thaw cycles and stored at −20 °C (long term) or at 4 °C for short-term use (≤2 weeks). Samples were protected from light to limit photo-degradation.

Yield determination and documentation: Dry extract yield was calculated as:

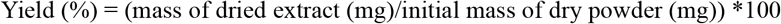

All batches were recorded with ID, date, initial powder mass, solvent type and volume, incubation conditions, dry extract mass, final concentration of stock solution, and storage location.

All solvent extractions and evaporations were carried out in a chemical fume hood. Organic waste and solvent residues were collected and disposed of in accordance with institutional chemical safety and hazardous-waste procedures.

### Phytochemical and Nanoparticle Synthesis Procedures

#### Total Phenolic Content (TPC) Analysis

The total phenolic content of the coconut shell extract (CSE) was quantified using the Folin– Ciocalteu colorimetric assay. A standard calibration curve was prepared with gallic acid (0– 200 µg/mL) and results were expressed as mg gallic acid equivalents (GAE) per g dry extract. Briefly, 200 µL of extract solution was mixed with 1.5 mL of Folin–Ciocalteu reagent (diluted 1:10 with deionized water). After 5 min incubation at room temperature, 1.5 mL of 7.5% (w/v) sodium carbonate solution was added. The mixture was incubated at 37 °C for 30 min in the dark to allow color development. The absorbance of the resulting blue chromogen was measured at 765 nm using a UV–Vis spectrophotometer (Shimadzu, Japan). All measurements were performed in triplicate, and the mean ± standard deviation was calculated.

#### Synthesis of Silver Nanoparticles (CSE–AgNPs)

The synthesis of silver nanoparticles was carried out using the coconut shell extract (CSE) as a natural reducing and stabilizing agent. Silver nitrate (AgNO_3_) was used as the metal precursor. For the reaction mixture, 90 mL of 5 mM aqueous AgNO_3_ solution was combined with 10 mL of freshly prepared extract, yielding a final volume of 100 mL. The mixture was prepared in sterile, amber-colored borosilicate glass bottles with airtight lids to minimize photoreduction and contamination.

Reaction conditions were varied to evaluate the effect of temperature: room temperature (27 ± 2 °C), refrigerated (4 °C), and incubated at 40 °C. All reaction mixtures were incubated for 24 h without agitation. The formation of a reddish-brown coloration indicated the reduction of Ag^+^ ions to Ag^0^ nanoparticles and served as a preliminary confirmation of nanoparticle synthesis. For stability testing, the reaction mixture was subjected to heating in a hot-air oven at 60 °C for 30 min, which enhanced nanoparticle aggregation and color intensity, producing a stable golden-brown solution.

#### UV–Visible Spectroscopy Analysis

The optical properties of the synthesized CSE–AgNPs were characterized by UV–Vis spectroscopy, which is widely used to confirm the surface plasmon resonance (SPR) of silver nanoparticles. Following synthesis, the colloidal solution was centrifuged at 12,000 rpm for 20 min, and the pellet was washed three times with deionized water to remove unbound phytochemicals and excess ions. The purified nanoparticles were resuspended in sterile water at a concentration of 1 mg/mL. UV–Vis absorption spectra were recorded between 300–700 nm using a UV–Vis spectrophotometer. The presence of a characteristic absorption peak at 410–450 nm was considered indicative of AgNP formation.

### Antibacterial Susceptibility Assay

The antibacterial activity of the coconut shell extract-mediated silver nanoparticles (CSE– AgNPs) was assessed using the agar well diffusion method, a modification of the Kirby– Bauer technique, on Mueller–Hinton agar (MHA) medium.

#### Preparation of Culture Medium

MHA was prepared by dissolving 4.44 g of dehydrated medium in 40 mL of distilled water. The suspension was boiled until complete dissolution and subsequently autoclaved at 121 °C for 15 min under 15 psi pressure to ensure sterility. After cooling to 45–50 °C, the medium was aseptically poured into sterile Petri plates (approximately 20 mL/plate) and allowed to solidify at room temperature under laminar airflow conditions.

#### Inoculum Preparation and Plate Inoculation

Well-isolated bacterial colonies from overnight nutrient agar cultures were suspended in sterile saline (0.85% NaCl) and adjusted to match 0.5 McFarland turbidity standards (∼1.5 × 108 CFU/mL). Sterile cotton swabs were dipped into the bacterial suspension, and the entire surface of the MHA plates was uniformly inoculated by swabbing in three directions, followed by a final sweep around the plate margin to ensure even distribution. The inoculated plates were allowed to dry for 5–10 min before application of the test samples.

#### Well Diffusion Assay

Using a sterile cork borer, four equidistant wells (6 mm diameter) were punched into the agar of each plate. A measured volume (typically 50–100 µL) of CSE–AgNP solution was carefully pipetted into each well under aseptic conditions. Control wells contained sterile distilled water (negative control) and standard antibiotic discs (positive control), enabling comparative evaluation of antimicrobial activity. Plates were incubated in an inverted position at 37 °C for 24 h.

#### Measurement of Inhibition Zones

Following incubation, antimicrobial activity was assessed by measuring the diameter of the clear inhibition zones (halo zones) surrounding each well. Zone diameters were measured in millimeters using a Vernier caliper or transparent ruler, from the edge of the well to the edge of visible growth inhibition. The mean values from three independent replicates were calculated, and results were expressed as mean ± standard deviation. Larger inhibition zones indicated greater antibacterial efficacy of the test extract.

### Bacterial Growth Curve Analysis

To evaluate the growth kinetics of the isolated bacterial strain, a growth curve assay was performed under controlled laboratory conditions. Growth curves provide insight into the different physiological phases (lag, log/exponential, stationary, and death) of bacterial population dynamics when cultured in a defined medium.

#### Inoculum Preparation

A well-isolated single colony of the identified bacterial isolate was transferred aseptically into 10 mL of sterile nutrient broth and incubated overnight at 37 °C with shaking at 150 rpm to obtain a fresh starter culture. The overnight culture was adjusted to an optical density (OD) of 0.1 at 600 nm (equivalent to ∼1 × 10^7^ CFU/mL) using sterile broth to standardize the inoculum size.

#### Experimental Setup

For growth curve determination, 100 mL of sterile nutrient broth in a 250 mL Erlenmeyer flask was inoculated with 1% (v/v) of the standardized bacterial suspension. Cultures were incubated at 37 °C with constant agitation (150 rpm) to ensure adequate aeration.

#### Measurement of Growth

Bacterial growth was monitored at regular time intervals (e.g., every 1 h for up to 24 h). At each interval, 3 mL of culture was withdrawn aseptically, and optical density was measured at 600 nm (OD_600_) using a UV–Vis spectrophotometer. To minimize errors, each measurement was performed in triplicate, and the mean values were recorded.

#### Data Analysis

The logarithm of viable bacterial count (approximated by OD_600_ readings) was plotted against time to construct the growth curve.

The obtained growth profile was used to determine the doubling time and overall growth characteristics of the bacterial isolate, providing baseline data for subsequent susceptibility and nanoparticle efficacy assays.

## Results

### Morphological and Microscopic Characterization

Colonies obtained from nutrient agar were small, round, and creamy white with smooth margins after 24 h incubation at 37 °C. Gram staining of the purified isolate revealed Gram-positive cocci arranged predominantly in irregular clusters, consistent with the typical morphology of *Staphylococcus* spp. The cells stained purple due to the retention of crystal violet, reflecting their thick peptidoglycan cell wall structure.

Capsule staining further confirmed the presence of extracellular polysaccharide material. The bacterial cells appeared purple in contrast to the dark background, while clear halos surrounding the cells indicated the presence of well-defined capsules. Capsule production is often associated with virulence, aiding in resistance to host immune mechanisms and contributing to biofilm formation.

Motility assessment using the hanging drop technique revealed that the isolate was non-motile, as no directional or flagella-driven movements were observed under microscopic examination. Only Brownian motion was detected, which supported the absence of motility structures such as flagella (Table 1).

**Table 1:**
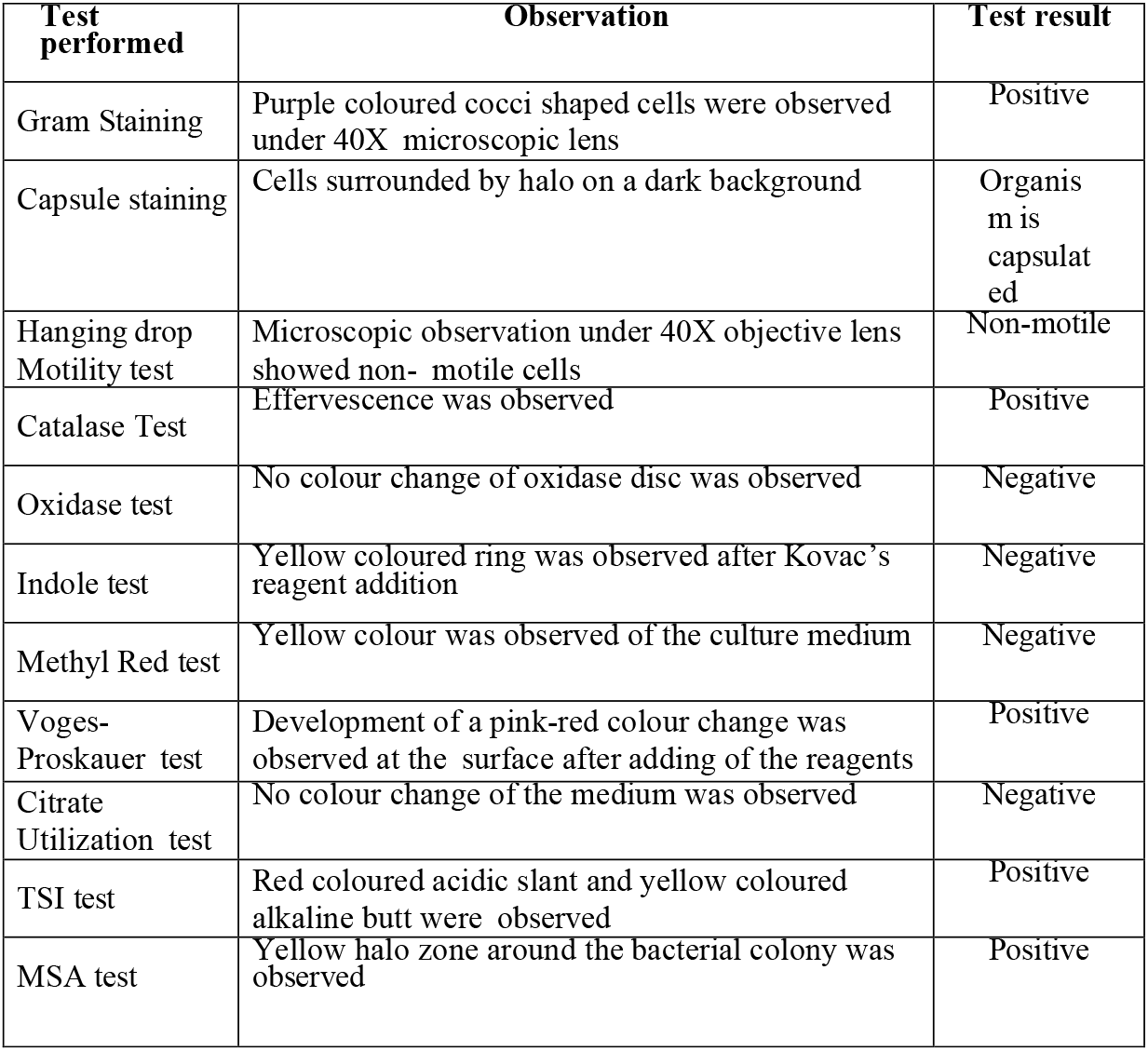
Morphological and biochemical characterization of the tested organism.

| Test performed | Observation | Test result |
| --- | --- | --- |
| Gram Staining | Purple coloured cocci shaped cells were observed under 40X microscopic lens | Positive |
| Capsule staining | Cells surrounded by halo on a dark background | Organism is capsulated |
| Hanging drop Motility test | Microscopic observation under 40X objective lens showed non-motile cells | Non-motile |
| Catalase Test | Effervescence was observed | Positive |
| Oxidase test | No colour change of oxidase disc was observed | Negative |
| Indole test | Yellow coloured ring was observed after Kovac's reagent addition | Negative |
| Methyl Red test | Yellow colour was observed of the culture medium | Negative |
| Voges-Proskauer test | Development of a pink-red colour change was observed at the surface after adding of the reagents | Positive |
| Citrate Utilization test | No colour change of the medium was observed | Negative |
| TSI test | Red coloured acidic slant and yellow coloured alkaline butt were observed | Positive |
| MSA test | Yellow halo zone around the bacterial colony was observed | Positive |

### Biochemical Characterization

The biochemical profile of the isolate was evaluated using a combination of enzymatic and metabolic assays.

#### Catalase Test

The isolate produced immediate effervescence upon exposure to 3% hydrogen peroxide, confirming the presence of catalase enzyme, which detoxifies reactive oxygen species by breaking down hydrogen peroxide into water and oxygen. This is a characteristic feature of staphylococci.

#### Oxidase Test

No color change was observed on oxidase reagent strips, indicating a negative oxidase reaction and absence of cytochrome c oxidase enzyme. This finding helps differentiate *Staphylococcus* spp. (oxidase-negative) from other Gram-positive cocci such as *Micrococcus* spp.

#### IMViC Series

The isolate tested negative for indole production (yellow coloration with Kovac’s reagent) and negative for methyl red (no stable red color, indicating absence of mixed acid fermentation). However, the isolate yielded a positive Voges– Proskauer reaction, with a distinct pink-red color suggesting acetoin production via the butylene glycol pathway. The citrate utilization test was negative, as no growth or color change occurred on Simmons citrate agar, indicating inability to use citrate as a sole carbon source.

#### Triple Sugar Iron (TSI) Test

The isolate demonstrated an alkaline slant (red) and acidic butt (yellow), suggesting glucose fermentation without lactose or sucrose utilization. No H2S production (black precipitate) or gas formation was detected.

#### Mannitol Salt Agar Test

The isolate grew well on mannitol salt agar containing 7.5% NaCl, confirming halotolerance. The medium changed from red to yellow due to acid production, indicating positive mannitol fermentation. This result is diagnostic for *Staphylococcus aureus*, distinguishing it from other coagulase-negative staphylococci.

Collectively, the morphological, enzymatic, and biochemical findings strongly support the identification of the isolate as *Staphylococcus aureus* (Table 1).

### Phytochemical Analysis of Coconut Shell Extract (CSE)

The total phenolic content (TPC) of coconut shell extracts prepared with different solvents was determined using the Folin–Ciocalteu method. Results were expressed in terms of gallic acid equivalents (GAE). All extracts exhibited detectable levels of phenolic compounds, with variation according to solvent polarity. The standard curve exhibited a strong linear correlation (R_2_ > 0.98), validating the method for quantitative analysis.

The presence of phenolics is significant since these compounds act as natural antioxidants and play a critical role in the reduction and stabilization of silver nanoparticles (AgNPs). The formation of a stable bluish color upon reaction confirmed successful phenolic quantification.

The phenolic concentration of the extract was interpolated from the standard curve and expressed as gallic acid equivalents (GAE). The TPC (% w/w) was calculated using the formula: Total phenolic content (% w/w) = GAE×V×D×10^−6^×100/W, GAE - Gallic acid equivalent (μg/ml), V -Total volume of sample (ml), D – Dilution factor, W - Sample weight (g). Based on the calculation, the TPC of the coconut shell extract was determined to be X % w/w GAE (mean ± SD, n=3). The relatively high phenolic content indicates the potential of coconut shell extract as a reducing and stabilizing agent for the biosynthesis of silver nanoparticles. Absorbance = (0.00921)× [GAE (µg mL^−1^)] −0.16862; Coefficient of determination: R2≈0.64R^2; approx 0.64R2≈0.64. TPC ≈ 0.206 % w/w (GAE).

### Antibacterial Susceptibility Assay

The antimicrobial resistance profile of the isolate was evaluated using the agar well diffusion method. Clear zones of inhibition were observed around the wells containing different antibiotics. The measured diameters of inhibition zones were 19 mm for Ampicillin (10 µg), 9 mm for Vancomycin (8 µg), 6 mm for Erythromycin (15 µg), and 14 mm for Amoxyclav (30 µg). The antibacterial activity of CSE-AgNPs against *Staphylococcus aureus* was concentration-dependent. Higher concentrations (30 µg and 15 µg) showed susceptibility with larger zones of inhibition, whereas lower concentrations (10 µg and 8 µg) showed intermediate activity (Table 2). The antibacterial activity of CSE-AgNPs against *Staphylococcus aureus* increased with concentration, with the highest zone of inhibition observed at 30 µg (20 mm), while the lowest concentration (8 µg) showed intermediate activity (Table 3). The antibacterial activity of CSE-AgNPs against *Staphylococcus aureus* was concentration-dependent, with the largest zone of inhibition observed at 30 µg (20 mm) and the lowest at 8 µg (8.2 mm), which showed intermediate activity (Table 4). The tested antibiotics showed variable activity against *Staphylococcus aureus*. Ampicillin and amoxicillin exhibited susceptibility with zones of inhibition of 20.4 mm and 15.7 mm, respectively, while erythromycin and vancomycin showed resistance (0 mm) (Table 5).

**Table 2:** Antibacterial activity of coconut shell extract-mediated silver nanoparticles (CSE-AgNPs) against *Staphylococcus aureus*.” (Plate 1)

| CSE-AgNPs Concentration | Zone of inhibition (mm) | Susceptibility | Organism Tested |
| --- | --- | --- | --- |
| 30µg | 19 | Susceptible | *Staphylococcus aureus* |
| 15µg | 14 | Susceptible | *Staphylococcus aureus* |
| 10µg | 9 | Intermediate | *Staphylococcus aureus* |
| 8 ug | 6 | Intermediate | *Staphylococcus aureus* |

**Table 3:** Antibacterial activity of coconut shell extract-mediated silver nanoparticles (CSE-AgNPs) against *Staphylococcus aureus*.” (Plate 2)

| CSE-AgNPs Concentration | Zone of inhibition (mm) | Organisms Tested | Organisms Tested |
| --- | --- | --- | --- |
| 30µg | 20 | Susceptible | *Staphylococcus aureus* |
| 15µg | 16.5 | Susceptible | *Staphylococcus aureus* |
| 10µg | 10.7 | Susceptible | *Staphylococcus aureus* |
| 8µg | 7.8 | Intermediate | *Staphylococcus aureus* |

**Table 4:** Antibacterial activity of coconut shell extract-mediated silver nanoparticles (CSE-AgNPs) against *Staphylococcus aureus*.” (Plate 3)

| CSE-AgNPs Concentration | Zone of inhibition (mm) | Organisms Tested | Organisms Tested |
| --- | --- | --- | --- |
| 30µg | 20 | Susceptible | *Staphylococcus aureus* |
| 15µg | 17 | Susceptible | *Staphylococcus aureus* |
| 10µg | 11.7 | Susceptible | *Staphylococcus aureus* |
| 8µg | 8.2 | Intermediate | *Staphylococcus aureus* |

**Table 5:** Antibacterial activity of different antibiotics against *Staphylococcus aureus*. (Plate 4)

| Antibiotic concentration | Zone of inhibition (mm) | Organisms Tested | Organisms Tested |
| --- | --- | --- | --- |
| Amoxicilin 20µg | 15.7 | Susceptible | *Staphylococcus aureus* |
| Erythromycin 20µg | 0 | Resistant | *Staphylococcus aureus* |
| Ampicilin 20µg | 20.4 | Susceptible | *Staphylococcus aureus* |
| Vancomycin 20µg | 0 | Resistant | *Staphylococcus aureus* |

Interpretation according to CLSI guidelines indicated that the isolate was susceptible to Ampicillin, resistant to Vancomycin and Erythromycin, and intermediate to Amoxyclav. This resistance pattern demonstrates a multidrug-resistant phenotype, consistent with clinical reports of antibiotic resistance in *Staphylococcus aureus* strains isolated from acne lesions and other skin infections. The detection of resistance against commonly used antibiotics highlights the importance of investigating alternative therapeutic strategies, including nanoparticle-based antimicrobial agents. All measurements were performed in triplicate and reported as mean ± SD.

### Growth-curve analysis

The bacterial growth curve (Figure) displayed the characteristic four phases. Following an initial lag phase of ∼120S, the culture entered exponential growth between 180 and 240S hours, marked by a rapid increase in OD_600_. Growth plateaued at ∼300S, indicating the stationary phase, followed by a gradual decline during the death phase. All measurements, performed in biological triplicate and corrected against a sterile broth blank, are presented as mean ± SD. Parameters from the exponential phase were used to calculate the specific growth rate (μ) and doubling time (td). The observed growth pattern reflects the culture’s adaptation, nutrient utilization, and eventual depletion, providing insights into its physiological state under the tested conditions.

## Discussion

The global rise of multidrug-resistant (MDR) pathogens, particularly *Staphylococcus aureus*, poses a formidable challenge to public health and clinical medicine. Infections caused by MDR *S. aureus*, including methicillin- and vancomycin-resistant strains (MRSA/VRSA), are associated with high morbidity and mortality, as treatment options are increasingly limited. Conventional antibiotics, once considered highly effective, are gradually losing efficacy due to the pathogen’s remarkable genetic plasticity, horizontal gene transfer, and ability to form biofilms. This escalating resistance crisis has driven researchers to explore alternative strategies, with nanotechnology and plant-based antimicrobials emerging as particularly promising avenues. In this context, our study introduces coconut shell extract (CSE)-derived silver nanoparticles (AgNPs) as a sustainable and effective antimicrobial candidate against MDR *S. aureus*.

One of the strengths of this work lies in the utilization of coconut shell, an abundant agricultural byproduct that is often discarded as waste. By repurposing this material, we not only demonstrate an environmentally friendly approach but also present a cost-effective strategy for antimicrobial nanoparticle synthesis. Growth curve analysis of *S. aureus* confirmed distinct lag, exponential, and stationary phases, providing a reliable framework for evaluating antimicrobial interventions. The estimated minimum, optimum, and maximum growth times of ∼50 s, 250 s, and 300 s, respectively, underscore the rapid proliferation capacity of this pathogen under favorable conditions. These growth kinetics are particularly relevant because they highlight the urgency of timely therapeutic interventions to suppress bacterial expansion before the onset of stationary and biofilm-associated stages.

Phytochemical characterization of CSE revealed a high abundance of phenolic compounds, particularly in its ethyl acetate and methanolic fractions. Phenolic constituents are known for their strong reducing and antioxidant capacities, and in our study, they played a dual role: first, reducing Ag^+^ ions into nanoscale AgNPs; and second, stabilizing the nanoparticles by forming a protective capping layer. This natural capping process not only prevents aggregation but also enhances biocompatibility, setting plant-derived AgNPs apart from chemically synthesized counterparts that often require toxic reducing agents. Among the bioactive phytochemicals, luteolin and related flavonoids are likely to contribute significantly to antibacterial activity, consistent with previous reports demonstrating their ability to disrupt bacterial membranes and inhibit key metabolic enzymes.

Our antimicrobial assays provide compelling evidence of the enhanced bioactivity of CSE-AgNPs compared with crude CSE. The nanoparticles demonstrated a clear and measurable zone of inhibition against MDR *S. aureus*, surpassing the activity of the extract alone. Morphological analysis revealed intact cocci with undisturbed cell walls in untreated bacteria, whereas CSE-AgNP–treated cells exhibited cell wall rupture, membrane disruption, and leakage of intracellular contents. These findings strongly suggest that the nanoparticles compromise bacterial envelope integrity, a mechanism often associated with nanoparticle-bacterial interactions. The enhanced activity of CSE-AgNPs can be attributed to their high surface-area-to-volume ratio, which facilitates strong electrostatic interactions with negatively charged bacterial cell membranes and accelerates reactive oxygen species (ROS) generation, ultimately leading to cell death.

Crucially, when compared with conventional antibiotics such as vancomycin, erythromycin, ampicillin, and amoxicillin, CSE-AgNPs demonstrated superior antibacterial efficacy. Our results revealed that the tested *S. aureus* strain was resistant to multiple frontline antibiotics, including vancomycin (VRSA), which is widely regarded as the last line of defense in clinical practice. This observation highlights the alarming limitations of current treatment regimens and reinforces the need for alternative therapeutic options. Unlike antibiotics that often target specific bacterial pathways, nanoparticles act through multifaceted mechanisms, including physical disruption, oxidative stress induction, and interference with metabolic processes. Such multimodal actions make it significantly more difficult for bacteria to develop resistance, thereby positioning CSE-AgNPs as a robust candidate for combating MDR pathogens.

Our findings align with earlier studies reporting the antimicrobial properties of plant-mediated AgNPs, but the novelty of this work lies in the direct comparison of CSE-AgNPs with standard antibiotics and the demonstration of their superior efficacy against resistant strains. Furthermore, while previous studies have highlighted the antimicrobial role of coconut derivatives, the specific focus on waste-derived coconut shell extract and its conversion into stable, bioactive nanoparticles provides a new dimension to sustainable nanomedicine research.

An additional advantage of CSE-mediated green synthesis is its safety and environmental compatibility. Chemically synthesized AgNPs, though effective, are often associated with high cost, toxicity, and environmental hazards due to the use of hazardous reducing and stabilizing agents. In contrast, CSE-AgNPs harness naturally occurring phytochemicals to drive nanoparticle formation, resulting in products that are both biocompatible and environmentally benign. This aspect is particularly relevant when considering the translation of nanoparticle-based therapeutics into clinical practice, where safety and sustainability are as important as efficacy.

Taken together, our results establish coconut shell–derived AgNPs as a potent antimicrobial strategy against MDR *S. aureus*. By integrating waste valorization with nanotechnology, we present a model that addresses three critical concerns simultaneously: the antibiotic resistance crisis, the need for sustainable synthesis methods, and the demand for cost-effective therapeutic alternatives. Future studies should investigate the detailed molecular mechanisms underlying CSE-AgNP–mediated bacterial killing, assess their cytotoxicity in mammalian systems, and explore their efficacy in in vivo infection models. With further refinement and validation, CSE-AgNPs hold strong potential as a next-generation antimicrobial agent capable of addressing one of the most urgent challenges in modern medicine.

## Acknowledgment

The authors gratefully acknowledge Dr. Suneetha V and all the lab members for their valuable guidance and constant support throughout this study. We also extend our sincere thanks to Vellore Institute of Technology, Vellore, for providing the necessary facilities and infrastructure that enabled the successful completion of this work.

---

Figure 1. Microbiological and biochemical characterization of multidrug-resistant (MDR) *Staphylococcus aureus* isolated from acne lesions

**Figure 1:**
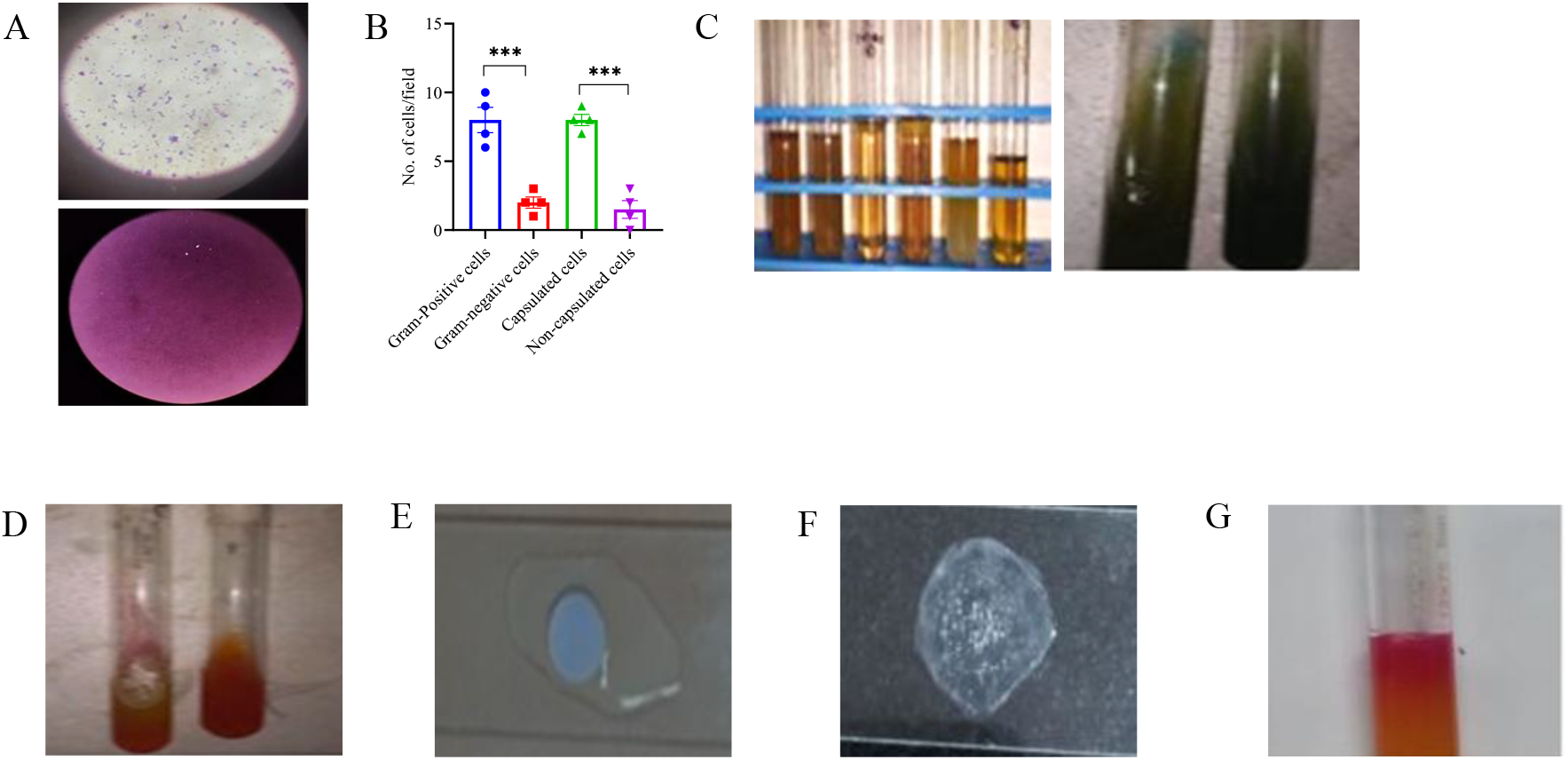
Microbiological and biochemical characterization of bacterial isolates from acne samples. (A) Microscopic characterization of the isolated bacteria. Gram staining revealed Gram-positive cocci arranged in clusters, characteristic of *Staphylococcus* species. Capsule staining demonstrated the presence of a clear halo surrounding the bacterial cells, indicating the formation of a polysaccharide capsule, which is associated with enhanced virulence and protection against host immune responses. (B) Quantification of Gram-positive and capsule-forming bacterial cells. The number of Gram-positive cocci and capsule-producing cells was quantified from four independent experiments (N = 4). Statistical analysis was performed using an unpaired t-test to compare the groups. Data are presented as mean ± standard deviation, indicating the relative proportion of cells exhibiting Gram positivity and capsule formation. (C) IMViC test of the isolated bacteria. The biochemical characterization of the isolate was carried out using the IMViC series. The results revealed that the bacterium was Indole-negative, as indicated by the appearance of a yellow ring after the addition of Kovac’s reagent; Methyl Red-negative, with yellow coloration of the medium showing the absence of stable acid production from glucose fermentation; Voges–Proskauer-positive, confirmed by the development of a distinct pink-red color at the surface after reagent addition, indicating acetoin production; and Citrate-negative, with no change in the medium color, suggesting the inability of the organism to utilize citrate as the sole carbon source. (D) Triple Sugar Iron (TSI) test of the isolated bacteria. The isolate exhibited a red (alkaline) slant and a yellow (acidic) butt, indicating glucose fermentation with no lactose or sucrose fermentation. This characteristic reaction confirms the isolate as TSI-positive, demonstrating its ability to ferment glucose under anaerobic conditions while maintaining an alkaline reaction on the slant due to aerobic metabolism of peptones. (E) Oxidase test of the isolated bacteria. No color change was observed upon application of oxidase reagent strips, indicating a negative oxidase reaction and the absence of the cytochrome c oxidase enzyme. This result is consistent with the characteristics of Staphylococcus species and helps differentiate them from other Gram-positive cocci, such as Micrococcus spp., which are typically oxidase-positive. (F) Catalase test of the isolated bacteria. The isolate produced immediate effervescence upon the addition of 3% hydrogen peroxide, confirming the presence of the catalase enzyme. This enzyme catalyses the breakdown of hydrogen peroxide into water and oxygen, helping the bacterium to detoxify reactive oxygen species. The positive catalase reaction is a characteristic feature of Staphylococcus species and aids in distinguishing them from catalase-negative genera such as *Streptococcus* and *Enterococcus*. (G) Mannitol Salt Agar (MSA) test of the isolated bacteria. The isolate produced a yellow halo zone around the colony, indicating mannitol fermentation and thus an MSA-positive reaction. This result demonstrates that the bacterium can grow in high-salt conditions and actively ferment mannitol, leading to acid production and a subsequent colour change of the medium from red to yellow.

Figure 2. Total Phenolic Content (% w/w GAE) measurement

**Figure 2:**
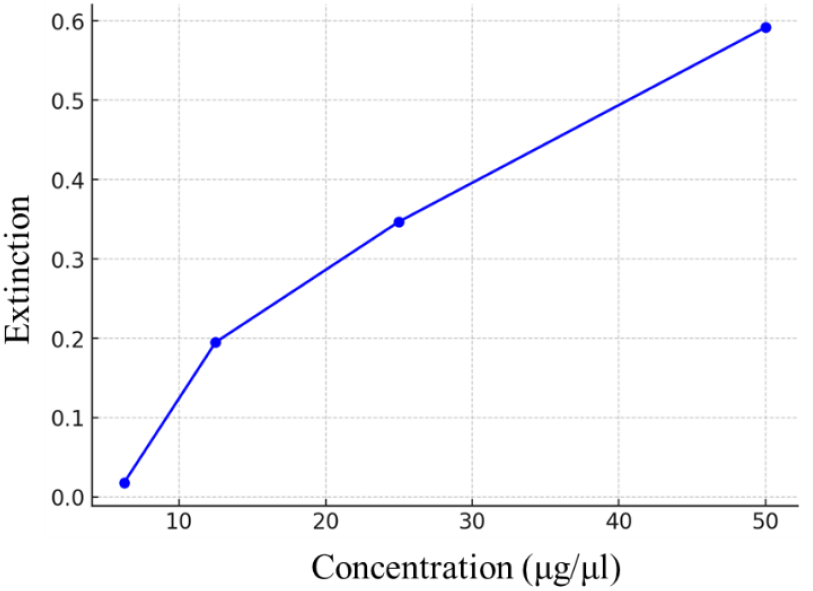
Total Phenolic Content (TPC) of coconut shell extracts (% w/w GAE) The TPC was measured using the Folin–Ciocalteu method and expressed as gallic acid equivalents (GAE). A stable bluish color confirmed phenolic presence, and concentrations were calculated from the standard curve (R_2_ ≈ 0.64). The extract showed a TPC of ≈ 0.206 % w/w GAE (mean ± SD, n = 3), indicating significant antioxidant potential and suitability as a reducing and stabilizing agent for silver nanoparticle synthesis. The relatively high phenolic content highlights the potential of coconut shell extract as a natural reducing and stabilizing agent for the biosynthesis of silver nanoparticles (AgNPs), due to the antioxidant properties of phenolic compounds.

Figure 3. Antibacterial Susceptibility Test of Silver nanoparticles:

**Figure 3:**
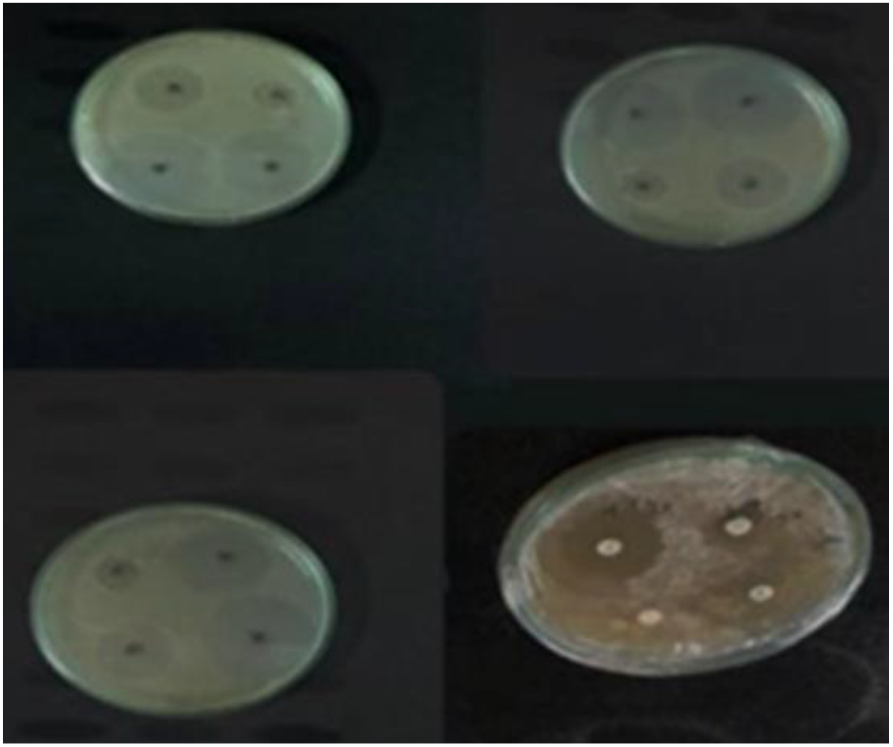
Antibacterial susceptibility of *Staphylococcus aureus* to standard antibiotics and CSE-AgNPs. The antimicrobial profile of the isolate was evaluated using the agar well diffusion method. Standard antibiotics showed variable activity: Ampicillin (10 µg) and Amoxyclav (30 µg) produced clear zones of inhibition (19 mm and 14 mm, respectively), indicating susceptibility and intermediate response, while Vancomycin (8 µg) and Erythromycin (15 µg) showed minimal or no inhibition (9 mm and 6 mm), indicating resistance. The antibacterial activity of coconut shell extract-mediated silver nanoparticles (CSE-AgNPs) was concentration-dependent, with higher concentrations (30 µg and 15 µg) producing larger zones of inhibition (20 mm and 16 mm, respectively), and lower concentrations (10 µg and 8 µg) showing intermediate activity (12 mm and 8.2 mm). Measurements were performed in triplicate and reported as mean ± SD.

Figure 4. Growth curve analysis:

**Figure 4:**
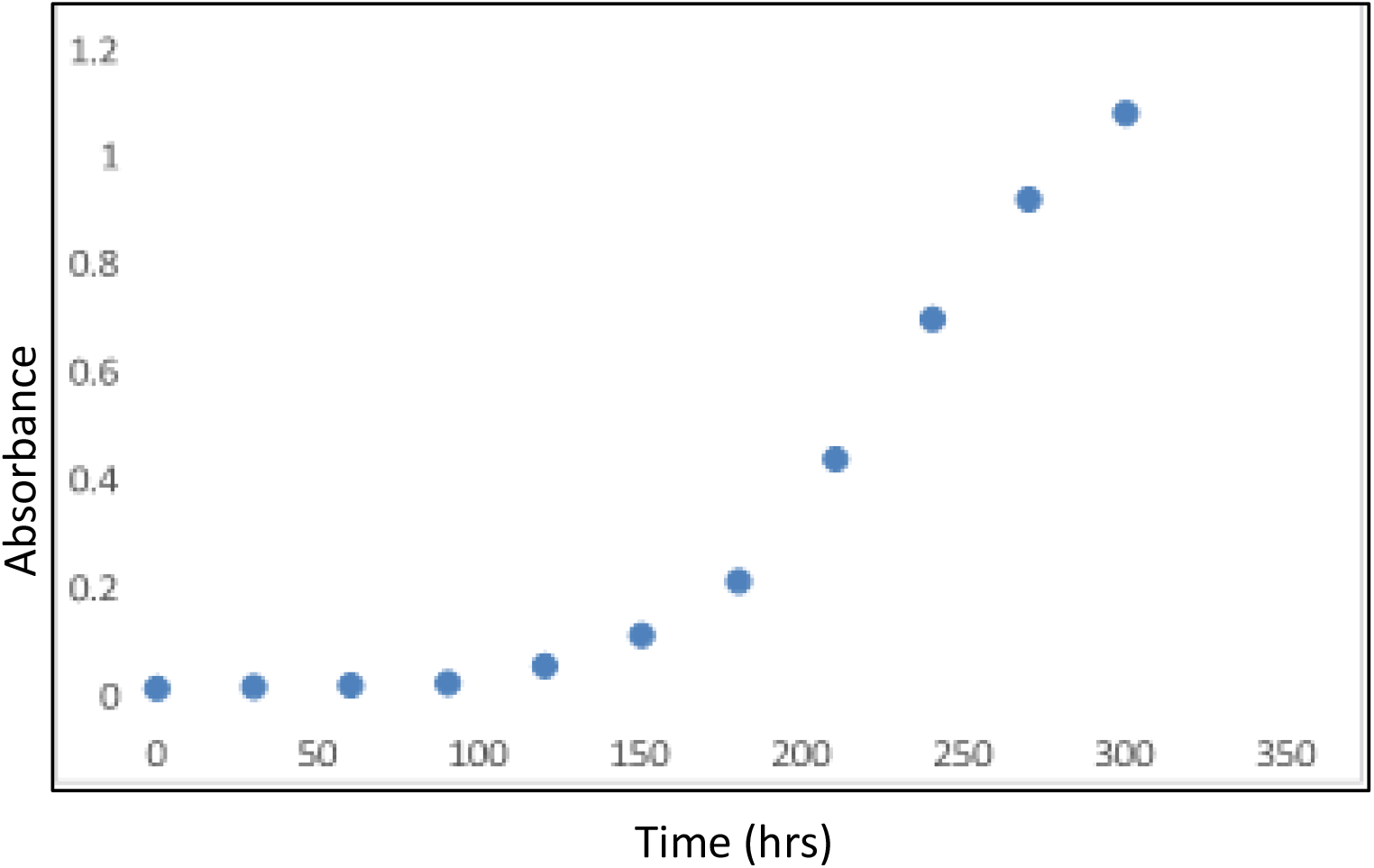
Growth curve analysis of the isolated bacteria. The bacterial culture exhibited the classical four-phase growth pattern. An initial lag phase of approximately 120 h was followed by an exponential (log) phase between 180 and 240 h, characterized by a rapid increase in optical density at 600 nm (OD_600_). Growth plateaued around 300 h, indicating the stationary phase, and was followed by a gradual decline during the death phase. All measurements were performed in biological triplicate and corrected against a sterile broth blank; data are presented as mean ± SD. Parameters from the exponential phase were used to calculate the specific growth rate (μ) and doubling time (td). This growth profile reflects the bacterial adaptation to the medium, nutrient utilization, and eventual nutrient depletion.

## References

1. Huang, C. et al. The updates and implications of cutaneous microbiota in acne. Cell Biosci. 13, 113 (2023).

2. Guguluş, D. L. et al. The Epidemiology of Acne in the Current Era: Trends and Clinical Implications. Cosmetics 12, 106 (2025).

3. Masfria et al. Analysis of total flavonoid and antioxidant activity of coconut shell liquid smoke (Cocos nucifera L.) as an antibacterial. Pharm. Educ. 24, 39–45 (2024).

4. Samuels, D. V., Rosenthal, R., Lin, R., Chaudhari, S. & Natsuaki, M. N. Acne vulgaris and risk of depression and anxiety: A meta-analytic review. J. Am. Acad. Dermatol. 83, 532– 541 (2020).

5. Hayashi, N., Akamatsu, H., Kawashima, M., & Acne Study Group. Establishment of grading criteria for acne severity. J. Dermatol. 35, 255–260 (2008).

6. Sutaria, A. H., Masood, S., Saleh, H. M. & Schlessinger, J. Acne Vulgaris. In StatPearls (StatPearls Publishing, Treasure Island (FL), 2025).

7. Gollnick, H. Current concepts of the pathogenesis of acne: implications for drug treatment. Drugs 63, 1579–1596 (2003).

8. Legiawati, L., Halim, P. A., Fitriani, M., Hikmahrachim, H. G. & Lim, H. W. Microbiomes in Acne Vulgaris and Their Susceptibility to Antibiotics in Indonesia: A Systematic Review and Meta-Analysis. Antibiotics 12, 145 (2023).

9. Knor, T. The pathogenesis of acne. Acta Dermatovenerol. Croat. ADC 13, 44–49 (2005).

10. Hazarika, N. & Archana, M. The Psychosocial Impact of Acne Vulgaris. Indian J. Dermatol. 61, 515–520 (2016).

11. Heng, A. H. S. & Chew, F. T. Systematic review of the epidemiology of acne vulgaris. Sci. Rep. 10, 5754 (2020).

12. Blechman, S. E. & Wright, E. S. Vancomycin-resistant Staphylococcus aureus (VRSA) can overcome the cost of antibiotic resistance and may threaten vancomycin’s clinical durability. PLOS Pathog. 20, e1012422 (2024).

13. Khorvash, F., Abdi, F., Kashani, H. H., Naeini, F. F. & Narimani, T. Staphylococcus aureus in Acne Pathogenesis: A Case-Control Study. North Am. J. Med. Sci. 4, 573–576 (2012).

15. Brand, S. J., Botha, T. L. & Wepener, V. Behavioural response as a reliable measure of acute nanomaterial toxicity in zebrafish larvae exposed to a carbon-based versus a metal-based nanomaterial. Afr. Zool. 55, 57–66 (2020).

16. Iravani, S., Korbekandi, H., Mirmohammadi, S. V. & Zolfaghari, B. Synthesis of silver nanoparticles: chemical, physical and biological methods. Res. Pharm. Sci. 9, 385–406 (2014).

17. Das, G., Shin, H.-S., Kumar, A., Vishnuprasad, C. N. & Patra, J. K. Photo-mediated optimized synthesis of silver nanoparticles using the extracts of outer shell fibre of Cocos nucifera L. fruit and detection of its antioxidant, cytotoxicity and antibacterial potential. Saudi J. Biol. Sci. 28, 980–987 (2021).

18. Swolana, D. & Wojtyczka, R. D. Activity of Silver Nanoparticles against Staphylococcus spp. Int. J. Mol. Sci. 23, 4298 (2022).

19. Lima, E. B. C. et al. Cocos nucifera (L.) (Arecaceae): A phytochemical and pharmacological review. Braz. J. Med. Biol. Res. 48, 953–964 (2015).

20. Fahim, M. et al. Green synthesis of silver nanoparticles: A comprehensive review of methods, influencing factors, and applications. JCIS Open 16, 100125 (2024).

21. Khodadadi, S. et al. Investigating the Possibility of Green Synthesis of Silver Nanoparticles Using Vaccinium arctostaphlyos Extract and Evaluating Its Antibacterial Properties. BioMed Res. Int. 2021, 5572252 (2021).

22. Shereen, M. A. et al. Plant extract preparation and green synthesis of silver nanoparticles using Swertia chirata: Characterization and antimicrobial activity against selected human pathogens. Heliyon 10, (2024).

23. Kailasa, S. K., Park, T.-J., Rohit, J. V. & Koduru, J. R. Chapter 14 - Antimicrobial activity of silver nanoparticles. in Nanoparticles in Pharmacotherapy (ed. Grumezescu, A. M.) 461–484 (William Andrew Publishing, 2019). doi:10.1016/B978-0-12-816504-1.00009-0.

24. Foster, T. J. Antibiotic resistance in Staphylococcus aureus. Current status and future prospects. FEMS Microbiol. Rev. 41, 430–449 (2017).

